# Endothelial Cell Dysfunction and Retinal Arteriolar Narrowing in Young Adults With Elevated Blood Pressure

**DOI:** 10.1101/2024.07.12.603349

**Authors:** Cheryl M. J. Tan, Adam J. Lewandowski, Henner Hanssen, Lukas Streese, Odaro J. Huckstep, Roman Fischer, Maryam Alsharqi, Afifah Mohamed, Wilby Williamson, Paul Leeson, Mariane Bertagnolli

## Abstract

**Background:** Young-adults with endothelial cell dysfunction are more likely to develop elevated blood pressure. We tested the hypothesis that this relates to development of structural microvascular impairments by studying associations between circulating endothelial colony-forming cell (ECFC) dysfunction and microvascular markers, as well as identifying related endothelial molecular mechanisms.

**Methods:** Peripheral blood ECFCs were isolated from 32 subjects (53% men, 28±4 years old) using the Ficoll density gradient centrifugation method. Participants with blood pressure ≥120/80 mm Hg were included in the elevated blood pressure (BP) group, whereas ≤120/80 mm Hg were classed as normotensive. Retinal microvasculature was assessed by Static Retinal Vessel Analyzer (SVA-T).

**Results:** Subjects with elevated BP had impaired in vitro ECFC colony-forming growth, cell proliferation and angiogenesis assessed by tube formation potential. There was a graded inverse association between ECFC colony-forming capacity (days taken for ECFC colony growth) and retinal arteriolar diameter, as well as arteriolar/venular ratio. Proteomic analysis of ECFCs identified differences in extracellular matrix organization, blood coagulation, exocytosis and vesicle transport proteins in subjects with elevated blood pressure, revealing the adaptor protein GRB2 as a potential link between endothelial cell and microvascular abnormalities.

**Conclusions:** Endothelial cell dysfunction associates with retinal arteriolar narrowing in men and women with elevated blood pressure. Endothelial molecular mechanisms linked to reduced adaptive postnatal angiogenesis capacity, rather than vascular development, may contribute to early microvascular changes.

**HIGHLIGHTS:**

- Subjects with elevated blood pressure had impaired in vitro endothelial cell growth and angiogenesis in comparison to normotensive subjects.
- There was an association between impaired endothelial cell growth capacity and reduced retinal arteriolar diameter.
- Different endothelial proteome signatures were identified, revealing the adaptor protein GRB2 as a potential link between endothelial and microvascular abnormalities in subjects with elevated blood pressure.

**GRAPHICAL ABSTRACT:** 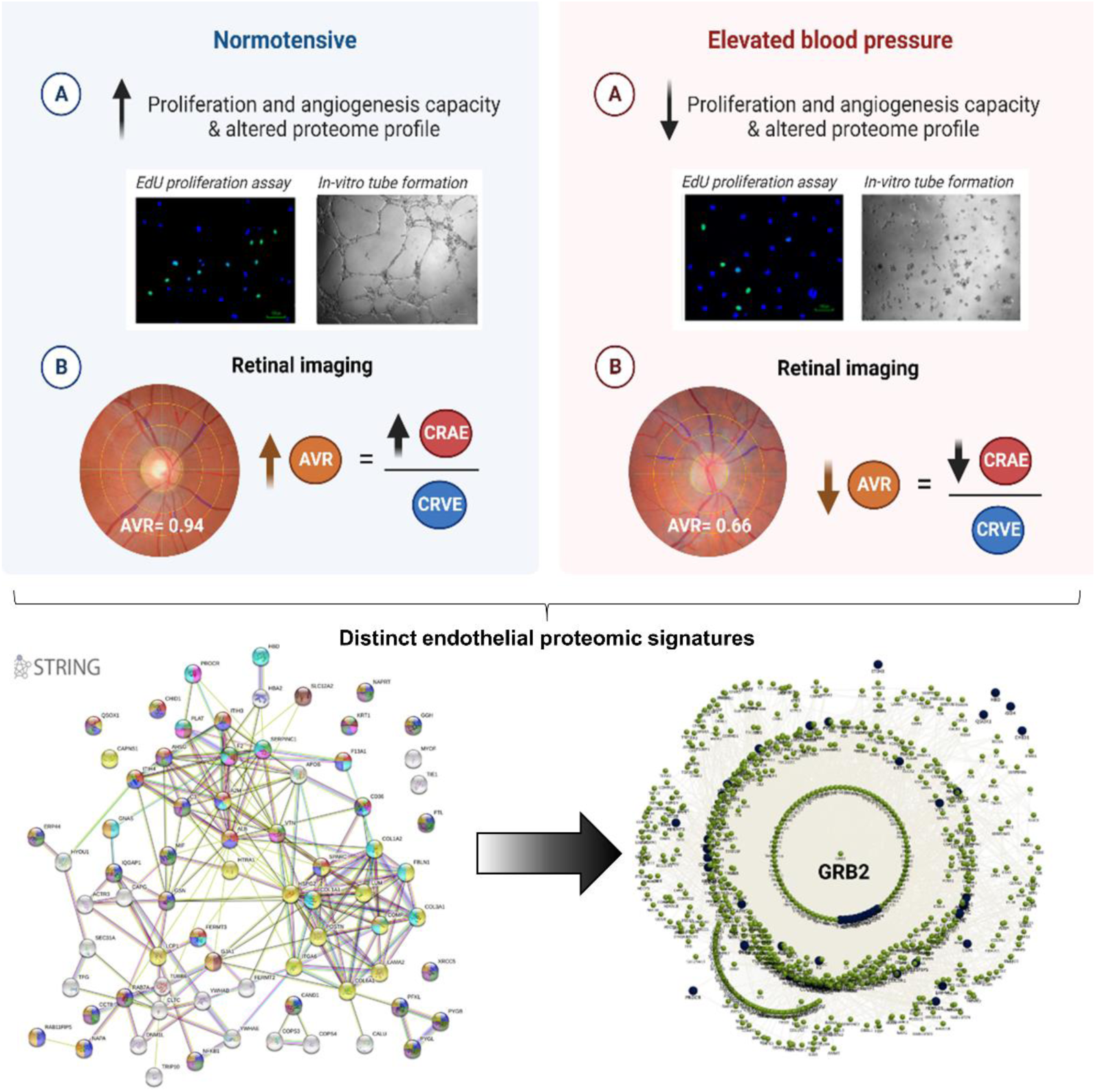

## INTRODUCTION

Endothelial cell and microcirculatory dysfunction have been observed in several cardiovascular diseases, including hypertension ^1–3^. Structural changes in microcirculation can relate to impaired angiogenic potential and reduced microvascular density or even with important structural changes characterized by narrower arterioles. Such alterations may increase peripheral vascular resistance, impair tissue perfusion and metabolism, and increase susceptibility to ischemia, therefore, playing an important role in the pathogenesis of hypertension ^4, 5^.

Endothelial and microcirculation alterations are caused by pro-thrombotic, pro-inflammatory, and/or pro-constrictive mechanisms in the arterioles, venules and capillaries ^4–7^. A growing number of studies have suggested the participation of endothelial cells in the pathogenesis of endothelial vascular dysfunction in hypertension ^2, 3, 8^. Recent investigations using circulating endothelial cells collected from patients’ peripheral blood have described an important association between circulating endothelial cell dysfunction and elevated blood pressure (BP) or with increased risk to develop hypertension across lifespan in young adult subjects ^2, 9, 10^. In most of these studies, endothelial colony-forming cells (ECFCs), a subpopulation of endothelial progenitor cells, were collected from peripheral blood, expanded in culture and functionally assessed *in vitro*, proposing a new method to clinically evaluate endothelial cell function in relationship with clinical characteristics. Despite the low number of ECFCs circulating in the blood, some clinical studies have shown that ECFC angiogenic and expansion capacities *in vitro* correlate with clinical characteristics such as BP ^2, 9, 11^ , or the risk of hypertension^2, 9^, pre-eclampsia^12^, diabetes^11, 13^ and bronchopulmonary dysplasia ^10^.

Adverse morphological alterations of the microvasculature are also seen during hypertension and corroborate to dysfunctional endothelial cells in these individuals ^1, 14^. Advances in fundus photography and image analysis techniques have enabled a highly reliable method to accurately visualize and measure retinal vessel diameters and function^15^ ^16–18^. Retinal microvasculature measurements include the diameters of retinal arterioles described as Central Retinal Arteriolar Equivalent (CRAE) and venules as Central Retinal Venular Equivalent (CRVE) ^5^. The microvascular architecture of retinal vessels has been increasingly linked to BP changes ^1, 19^ with retinal arteriolar narrowing in patients with elevated BP ^5, 19–21^. However, there is still an important knowledge gap on mechanisms driving these microvascular changes in young adults with elevated BP, who have a higher risk of developing hypertension.

We postulate that ECFC dysfunction can associate with microvascular impairments in young adult subjects with elevated BP. Therefore, in this study we aimed to assess correlations between circulating ECFC function and retinal microvascular structure in young adult subjects with elevated BP, and to identify endothelial proteomic signatures to target mechanisms linked to microvascular alterations.

## METHODS

### Study population

This study was designed and conducted in the Cardiovascular Clinical Research Facility, University of Oxford, John Radcliffe Hospital, United Kingdom. Participants were recruited as part of an open, parallel-arm randomised controlled trial - Trial of Exercise to Prevent HypeRtension in young Adults (TEPHRA) study (Ethics no. 16/SC/0016; Clinicaltrials.gov registration number NCT02723552. ^22, 23^ For this study, data were collected from 32 participants aged 18-35 years, BMI <30 kg/m^2^, clinical BP <159 mm Hg systolic, and/or 99 mm Hg diastolic BP without any clinical diagnosis of cardiovascular diseases and medications. All participants provided written consent for collecting and subsequent experimental use of samples in compliance with appropriate ethical approvals. Briefly, participants of the TEPHRA study underwent several comprehensive clinical measurements, including fasting blood, microvascular and cardiovascular assessments as previously described ^22^. Trained study investigators conducted all measurements.

### Anthropometry

Height and weight were measured to the nearest 0.1 cm and 0.1 kg using Seca measuring station (Seca, Birmingham, United Kingdom). Footwear was removed before the measurements. Manual waist circumference was measured with a tape at 2cm above the iliac crest, and the manual hip circumference was measured at the broadest portion near the level of the greater trochanter of the femur or mid-buttock. The waist and hip circumference ratio were then calculated.

### Blood pressure measurements

The brachial BP was measured using an automated oscillometer device monitor (Dinamap V100, GE Healthcare, Chalfont St. Giles, United Kingdom) after 5 minutes of resting in a seated position. Three readings were taken from the left arm, with the last two readings averaged and subsequently used for further analyses. Participants with BP >120/80 mm Hg were all categorised in the elevated BP group, whereas ≤120/80 mm Hg were classified as normotensive.

### Peak VO_2_/kg Measurement

Peak VO_2_/kg was measured through a standardised cardiopulmonary exercise test (CPET) conducted in a controlled laboratory environment as previously described. Briefly, participants performed a graded exercise test on a treadmill or cycle ergometer, with continuous monitoring of oxygen consumption (VO_2_) and carbon dioxide production (VCO_2_) using a metabolic cart. The highest VO_2_ value achieved during the test, adjusted for body weight (kg), was recorded as Peak VO_2_/kg.

### Physical Activity Monitoring

Participants were asked to wear an Axivity AX3 accelerometer continuously for seven days. The Axivity AX3 (Axivity Ltd, Newcastle Upon Tyne, United Kingdom), similar in design to a wrist-worn watch, has high compliance and reliability and is a validated measure of physical activity. Before the start of data collection, the Axivity was synchronized and initialized to start measuring simultaneously with the ActiGraph using the Open Movement software (OMGui; Newcastle University, UK). Data were processed using Open Movement software algorithm to categorize physical activity into different intensity levels – sedentary behaviour, light-intensity, walking, moderate, moderate-vigorous and vigorous physical activity. Vigorous physical activity was defined according to established metabolic equivalent (MET) thresholds. The total time spent in vigorous physical activity each day was summed across the week to calculate the average daily hours.

### Cardiovascular Fitness Assessment

Peak VO_2_/kg values were compared to the average daily hours of vigorous physical activity to assess their relationship with cardiovascular fitness levels. By integrating Peak VO_2_/kg measurements with accelerometer-based activity monitoring, a comprehensive assessment of participants’ cardiovascular fitness was achieved.

### Static retinal vessel analysis (SVA)

Static Retinal Vessel Analyzer (SVA-T, Imedos Systems, Germany) was used to capture three valid images of the retina of the left and the right eye, with the optic disk in the centre for static retinal vessel analysis. The detailed procedure was described previously ^24, 25^. Briefly, retinal arterioles and venules, coursing through an area of 0.5–1 disc diameter from the margin of the optic disc, were identified semi-automatically using an analyzing software identifying retinal vessels in ring-zones (Vesselmap 2, Visualis, Imedos Systems UG).

Diameters were averaged to CRAE and CRVE using the Parr-Hubbard formula ^26^. CRAE and CRVE values were used to calculate the AVR, taking the mean of the right and left eye results.

### Blood sampling

Participants were instructed to have fasted at least 4 hours prior to blood sample collection. All blood samples were collected from the antecubital fossa by venepuncture. A total volume of 50 mL of peripheral venous blood was collected per individual. Blood samples were centrifuged within 30 min, and the separated plasma and serum were stored at - 80°C for further analysis. Fasting blood biochemistry was measured at the Oxford John Radcliffe Hospital Biochemistry Laboratory using routine validated assays with clinical level quality controls. Insulin resistance was calculated using the homeostasis model assessment calculator ^27^.

### ECFC isolation and culture

The protocol of peripheral blood mononuclear cell (PBMC) isolation was previously described by us ^2, 10^. Firstly, 15 ml of fasted whole blood were diluted 1:1 with Dulbecco’s phosphate-buffered saline (PBS). PBMCs were separated from the diluted blood using 10 ml Ficoll-Paque PLUS (GE Healthcare Life Sciences, USA) density gradient following 30 min of 1,500 rpm centrifugation at room temperature and then washed twice with PBS (Gibco by Life technologies). PBMCs were either frozen down in 10% dimethyl sulfoxide (Sigma Aldrich) fetal bovine serum (FBS, ThermoFisher Scientific, Gibco, USA) or immediately plated on collagen I-coated (Corning, Corning, NY, USA) 25 cm^2^ tissue culture-treated Falcon flasks (ThermoFisher Scientific, USA) at a density of 5.0x10^6^ cells/flask. Cultured cells were maintained in a standard cell culture condition (humidified chamber, at 37°C with 21% O_2_ and 5% CO_2_), using complete endothelial cell growth basal medium-2 (EBM™-2 plus SingleQuots^TM^ of Growth Supplements, Lonza, Basel, Switzerland) supplemented with 1% penicillin/streptomycin (ThermoFisher Scientific, Gibco, USA) and 10% FBS.

Media of the cultured PBMCs was changed every two or three days, and cells were maintained for up to 30 days in culture. Cultured cells were observed daily from days six to 30 to determine the first day of cobblestone patterned ECFC colony formation. Once ECFC colony was observed, cells were passaged and expanded under the same conditions. ECFC functions were assessed using cells from the second passage. The remaining ECFCs were frozen and stored in standard conditions.

### In vitro tube formation assay

Vascular tube formation was assessed *in vitro* by plating 1.5x10^4^ cells on 50 µL of growth factor reduced basement membrane matrix (Matrigel) in a 96-well plate. Cells were imaged using a Leica DMIL inverted trinocular phase-contrast microscope (Leica, Germany) and 5x objective after 6 hours of incubation under standard culture conditions as illustrated in **Fig.1A**. Assays were performed in triplicates, and the number of closed tubes and branches formed (not necessarily forming closed tubes) were quantified in three to six random images per participant using ImageJ software (NIH, USA).

**Figure 1.**
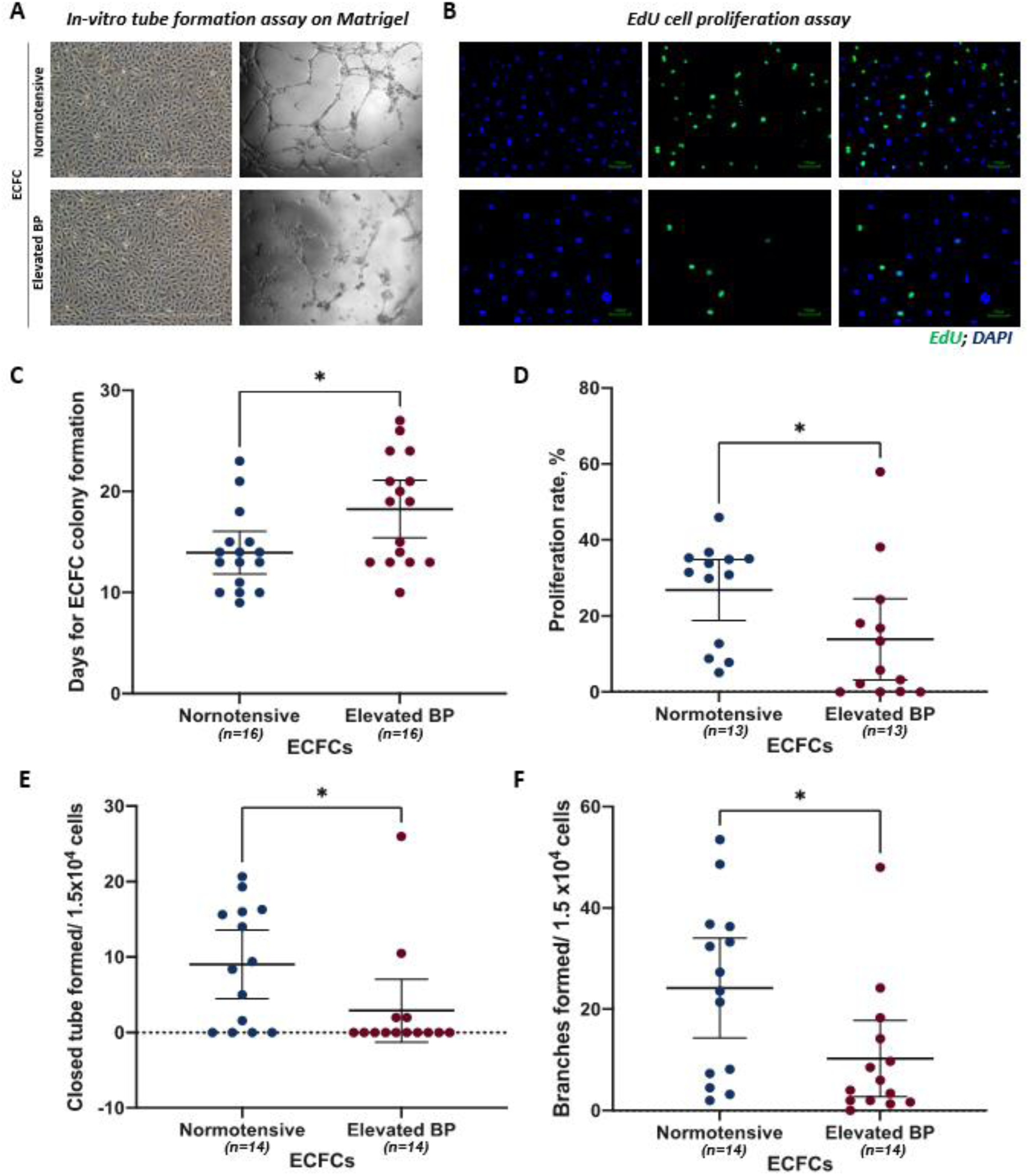
Comparisons of endothelial colony-forming cells (ECFCs) functional features between subjects with normal versus elevated blood pressure (BP). **A**, ECFC tube formation capacity was assessed using Matrigel tube formation assay with the numbers of total branches and closed tubes formed counted in phase-contrast images (5x objective). B, Proliferation rate was determined by DNA Edu incorporation (AlexaFluor 488, green), with nuclei counterstained with 4’,6-diamidino-2-phenylindole (DAPI, blue) (10x objective). C-F, Scatter plots representing respective ECFC clusters for each of the following cell characteristics: C, days for ECFC colony formation, D, proliferation rate (%), E, total closed tubes formed per 1.5x10^4^ cells, and F, total branching formed per 1.5x10^4^ cells. Comparison between normotensive and elevated BP groups by ANCOVA having sex as covariate. Data presented as mean ± standard deviation (SD). * P<0.05.

### ECFC proliferation assay

The proliferation rate was assessed by quantifying 5-ethynyl-2’-deoxyuridine (EdU) cellular incorporation using the Click-iTTM EdU Alexa FluorTM 488 Imaging kit (Thermo Fisher Scientific). A number of 4.0x10^4^ ECFCs were plated on collagen-coated flasks 1.86 cm^2^ surface area, maintained with complete EBM™-2 media and grown for 24 hours. Cells were then incubated with 10 µM EdU in complete EBM™-2 media for 4.5 hours at 37° C, then fixed with 3.7% formaldehyde in PBS, permeabilised with 0.5% Triton X-100 in PBS, and stained according to kit instructions. Assays were performed in triplicates, and cell images were obtained using a fluorescence microscope (Leica, Germany) and a 10x objective as illustrated in **Fig.1B**. The number of cells under proliferation and incorporating EdU into their DNA, stained positive for EdU (green, AlexaFluor 488), were counted in three to five full pictures per assay, as well as the total number of cells with nuclei stained with DAPI (blue); the percentage of proliferating cells (a measure of proliferation rate) was then calculated.

### Sample digestion and de-saltation

Age and sex-matched ECFCs from normotensive and elevated BP subjects (total n=8) were used for proteomic analysis at passage number two. Cell pellets containing approximately 2.0 x10^5^ were re-suspended in lysis buffer (50 mM Tris-HCl pH 8.5, 4% sodium dodecyl sulfate (SDS) and 50 mM dithiothreitol), boiled for 5 minutes and incubated for more than 30 min at room temperature for protein solubilisation. Total protein (∼250 µg) was digested and desalted after two rounds of chloroform-methanol precipitation, followed by reduction/alkylation with DTT/Iodoacetamide ^28^. Digested and desalted proteins were dissolved in 0.1% formic acid and loaded into the nano-LC-MS/MS system (Orbitrap Fusion Lumos, Thermo Fisher Scientific) in collaboration with the Oxford Target Discovery Institute.

### Mass Spectrometry report system and condition**s**

Mass spectrometry (MS) analysis was performed as described previously by us ^2^. Peptides were separated with a gradient of 2-35% acetonitrile in 0.1% formic acid/5% DMSO on an EasySpray column (50 cm x 75 µm) over 60 minutes and with a flow rate of 250 nl/min. MS data were acquired on an Orbitrap Fusion Lumos using a standard method as described previously ^29^.

### Proteomics data availability and system biology analyses

Peptides and proteins were identified by searching the MS raw files against the Human SwissProt database downloaded on 11/2015 (containing 20,268 human sequences). MASCOT data outputs were filtered by applying a 20-ion cutoff and 1% False Discovery Rate (FDR) above identity or homology threshold. The raw MS proteomics data have been deposited to the ProteomeXchange Consortium (http://www.proteomexchange.org) via the PRIDE partner repository with the dataset identifier: PXD020677 and project DOI: 10.6019/PXD020677.

Perseus (Max Planck Institute of Biochemistry, Germany)^30^ was used for quantitative analysis of the log-transformed (normalised protein abundance) after label-free quantitation as performed in Progenesis QI (Waters) using default settings. The data were filtered per row for 70% of the valid values, followed by imputation of missing values and ≥ 2 identified peptides. Student’s T-tests were used to make comparisons between the groups and genes satisfying a difference of -1.5 < Log2(fold change) >1.5, and FDR-corrected p-value (named q-value) <0.05 were considered statistically significant. As part of the proteomic analysis, all the p-values were expressed as FDR-corrected p-value (q-value) unless stated otherwise.

Gene Ontology (GO) enrichment was used with the following ontology sources: KEGG Pathway, GO Biological Processes, Reactome Gene Sets, Canonical Pathways, and CORUM using Metascape (http://www.metascape.org)^31^. Functional protein association networks of identified proteins were created using STRING (http://www.string-db.org) and InstantClue software ^32^. Gene Ontology enRIchement anaLysis and visuaLisAtion (Gorilla) (http://cbl-gorilla.cs.technion.ac.il/) was used for identifying and visualising enriched GO terms in ranked lists of genes ^33^.

### Statistical analysis

Statistical analysis was performed using SPSS (version 25, IBM, North Castle, NY, USA). Shapiro-Wilk testing and visual inspection (QQ-plots and histograms) were used to assess whether variables were normally distributed. Dependent on the normality of sample distribution, comparisons between normotensive and elevated BP groups were performed by either parametric independent t-test or non-parametric Mann-Whitney U test. Categorical variables were compared by χ2 or Fisher’s Exact tests. Comparisons between groups were performed by ANCOVA statistical test having only sex or sex and glucose as covariates.

Linear regression modelling was performed using forced entry with missing data excluded pairwise. Model analyses were performed using multiple linear regression with systolic BP being the dependent variable and 95% confidence intervals (CIs) reported for the variables of interest. Standardised regression coefficients (β) with 95% CI, standardised coefficients (β), and P values are reported. Results are presented as mean ± standard deviation (SD) unless stated otherwise. P values ≤ 0.05 were considered statistically significant.

## RESULTS

### Clinical characteristics of the study population

Clinical characteristics of our study population are presented in **Table 1**. No difference was observed between groups for the number of male/female subjects or their age, height, body mass, body mass index (BMI), and cardiovascular fitness. Significant differences were observed between groups in resting systolic (114±8 versus 133±10 mmHg, P<0.001), diastolic (69±7 versus 87±7 mm Hg, P<0.001), and mean BP (86±6 versus 104±7 mm Hg, P<0.001). Both groups displayed similar blood biochemistry profiles, with no significant differences observed in triglycerides, lipid profiles, insulin, Homeostasis Model Assessment (HOMA) β-cell, HOMA β-cell sensitivity and HOMA insulin resistance between groups apart from a higher fasting glucose levels in the elevated BP group (4.7±0.2 versus 5.1±0.4 mmol/L, P=0.01).

**Table 1.**
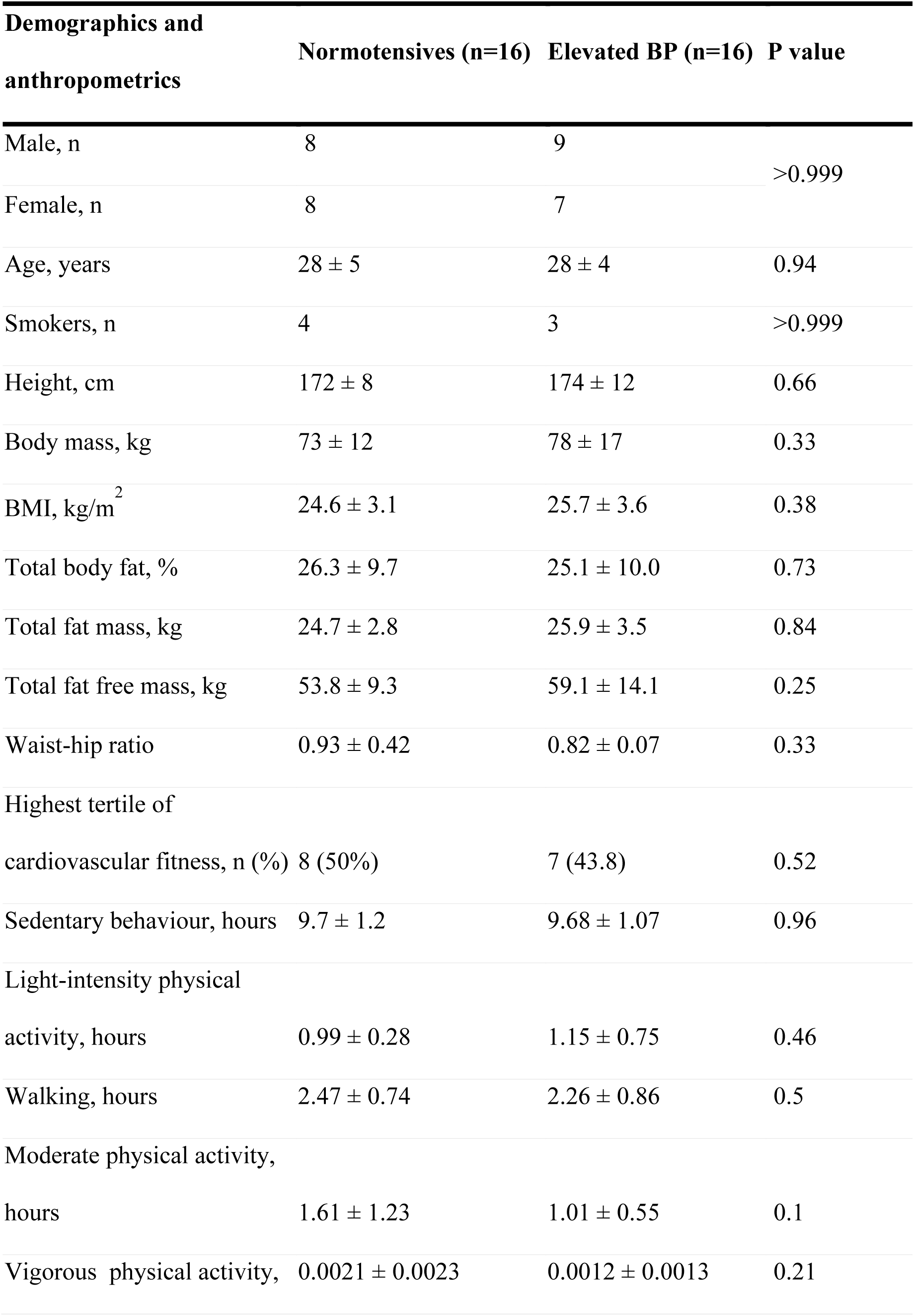
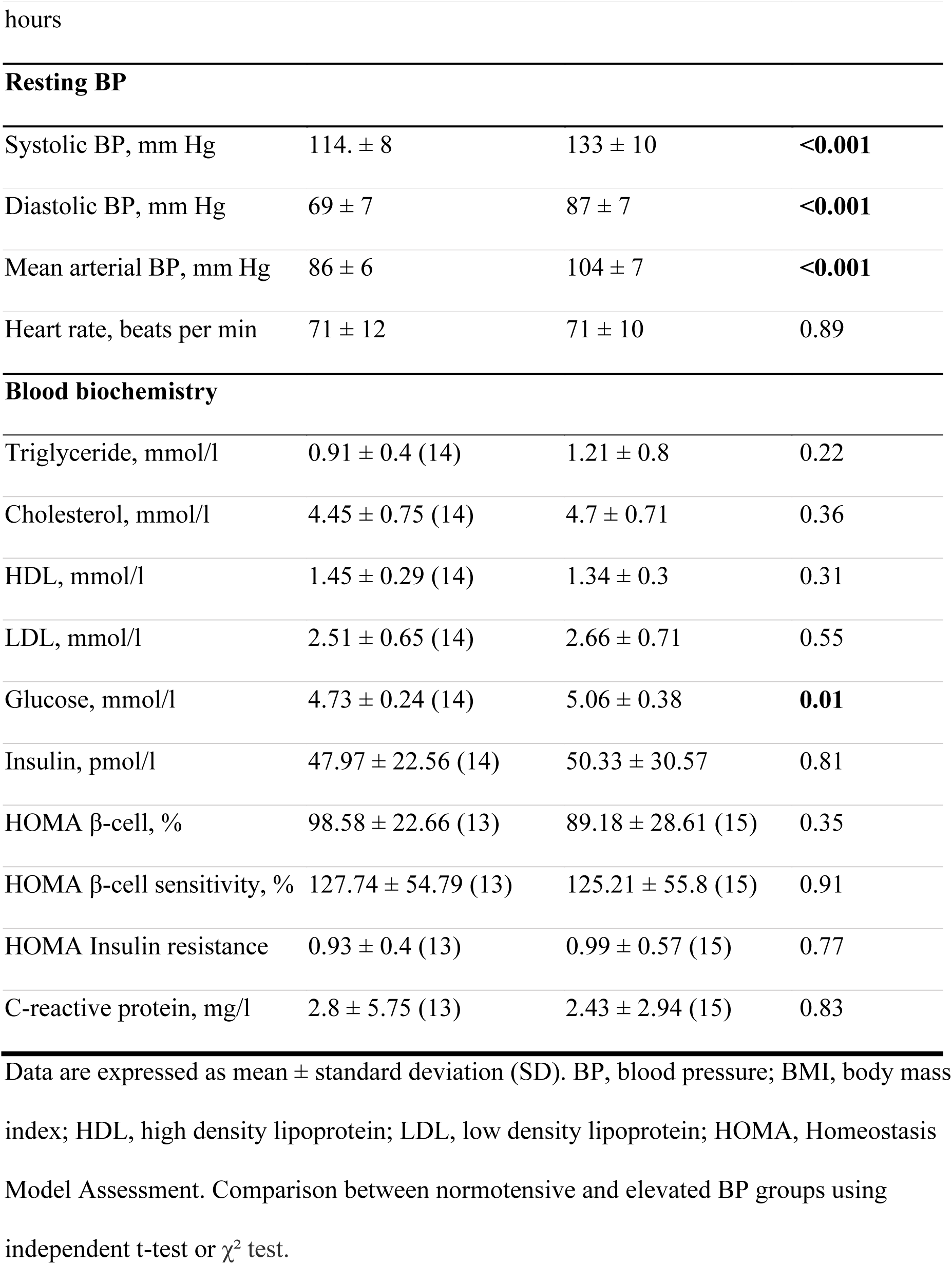
Clinical characteristics of the study population.

| **Demographics and anthropometrics** | **Normotensives (n=16)** | **Elevated BP (n=16)** | **P value** |
| --- | --- | --- | --- |
| Male, n | 8 | 9 | >0.999 |
| Female, n | 8 | 7 |  |
| Age, years | 28 ± 5 | 28 ± 4 | 0.94 |
| Smokers, n | 4 | 3 | >0.999 |
| Height, cm | 172 ± 8 | 174 ± 12 | 0.66 |
| Body mass, kg | 73 ± 12 | 78 ± 17 | 0.33 |
| BMI, kg/m<sup>2</sup> | 24.6 ± 3.1 | 25.7 ± 3.6 | 0.38 |
| Total body fat, % | 26.3 ± 9.7 | 25.1 ± 10.0 | 0.73 |
| Total fat mass, kg | 24.7 ± 2.8 | 25.9 ± 3.5 | 0.84 |
| Total fat free mass, kg | 53.8 ± 9.3 | 59.1 ± 14.1 | 0.25 |
| Waist-hip ratio | 0.93 ± 0.42 | 0.82 ± 0.07 | 0.33 |
| Highest tertile of cardiovascular fitness, n (%) | 8 (50%) | 7 (43.8) | 0.52 |
| Sedentary behaviour, hours | 9.7 ± 1.2 | 9.68 ± 1.07 | 0.96 |
| Light-intensity physical activity, hours | 0.99 ± 0.28 | 1.15 ± 0.75 | 0.46 |
| Walking, hours | 2.47 ± 0.74 | 2.26 ± 0.86 | 0.5 |
| Moderate physical activity, hours | 1.61 ± 1.23 | 1.01 ± 0.55 | 0.1 |
| Vigorous physical activity, hours | 0.0021 ± 0.0023 | 0.0012 ± 0.0013 | 0.21 |
| **Resting BP** |  |  |  |
| Systolic BP, mm Hg | 114. ± 8 | 133 ± 10 | **<0.001** |
| Diastolic BP, mm Hg | 69 ± 7 | 87 ± 7 | **<0.001** |
| Mean arterial BP, mm Hg | 86 ± 6 | 104 ± 7 | **<0.001** |
| Heart rate, beats per min | 71 ± 12 | 71 ± 10 | 0.89 |
| **Blood biochemistry** |  |  |  |
| Triglyceride, mmol/l | 0.91 ± 0.4 (14) | 1.21 ± 0.8 | 0.22 |
| Cholesterol, mmol/l | 4.45 ± 0.75 (14) | 4.7 ± 0.71 | 0.36 |
| HDL, mmol/l | 1.45 ± 0.29 (14) | 1.34 ± 0.3 | 0.31 |
| LDL, mmol/l | 2.51 ± 0.65 (14) | 2.66 ± 0.71 | 0.55 |
| Glucose, mmol/l | 4.73 ± 0.24 (14) | 5.06 ± 0.38 | **0.01** |
| Insulin, pmol/l | 47.97 ± 22.56 (14) | 50.33 ± 30.57 | 0.81 |
| HOMA β-cell, % | 98.58 ± 22.66 (13) | 89.18 ± 28.61 (15) | 0.35 |
| HOMA β-cell sensitivity, % | 127.74 ± 54.79 (13) | 125.21 ± 55.8 (15) | 0.91 |
| HOMA Insulin resistance | 0.93 ± 0.4 (13) | 0.99 ± 0.57 (15) | 0.77 |
| C-reactive protein, mg/l | 2.8 ± 5.75 (13) | 2.43 ± 2.94 (15) | 0.83 |

Data are expressed as mean ± standard deviation (SD). BP, blood pressure; BMI, body mass index; HDL, high density lipoprotein; LDL, low density lipoprotein; HOMA, Homeostasis Model Assessment. Comparison between normotensive and elevated BP groups using independent t-test or $\chi^2$ test.

### ECFC function in subjects with normal versus elevated BP

Comparisons of ECFC functional characteristics between groups adjusted by sex revealed that subjects with elevated BP have slower ECFC colony growth (18.3 ± 5.4 versus 13.9 ± 3.9 days, P=0.016, **Fig 1C**). Some ECFCs had grown small colonies in the first 30 days in culture but did not reach enough confluence to perform additional functional tests.

From those reaching a successful expansion, ECFCs of subjects with elevated BP had lower proliferation rate (13.9 ± 17.7 versus 26.8 ± 13.3 % of total cells, P=0.037, **Fig.1D**), and reduced angiogenic capacity indicated by the number of closed tubes formed *in vitro* (2.9 ± 7.2 versus 9.0 ± 7.9 number of closed tubes per 1.5 x 10^4^ cells, P=0.034, **Fig.1E**) and the number of tube branching (10.2 ± 13.0 versus 24.1 ± 17.1 number of branches per 1.5 x 10^4^ cells, P=0.017, **Fig.1F**) compared to normotensive subjects. However, after adjusting these comparisons for glucose levels, days to form ECFC (P=0.023) and proliferation rate (P=0.033) remained significantly different but numbers of closed tubes (P=0.092) and branches formed (P=0.059) were no longer significantly different.

### Retinal vessel diameters in subjects with normal versus elevated BP

Fundoscopy of both left and right eyes of the participants revealed that the group of subjects with elevated BP had lower CRAE (169.1 ± 20.0 μm vs 187.2 ± 11.2 μm, P<0.01, **Fig.2A**) indicating retinal arteriolar narrowing, without significant difference in CRVE values between groups (**Fig.2B**). Consequently, the arteriolar-to-venular-ratio (AVR) was lower in subjects with elevated BP (0.81 ± 0.07 vs. 0.88 ± 0.07, P<0.001, **Fig.2C**). Differences on CRAE and AVR remained statistically significant after adjustment for sex and glucose levels.

**Figure 2.**
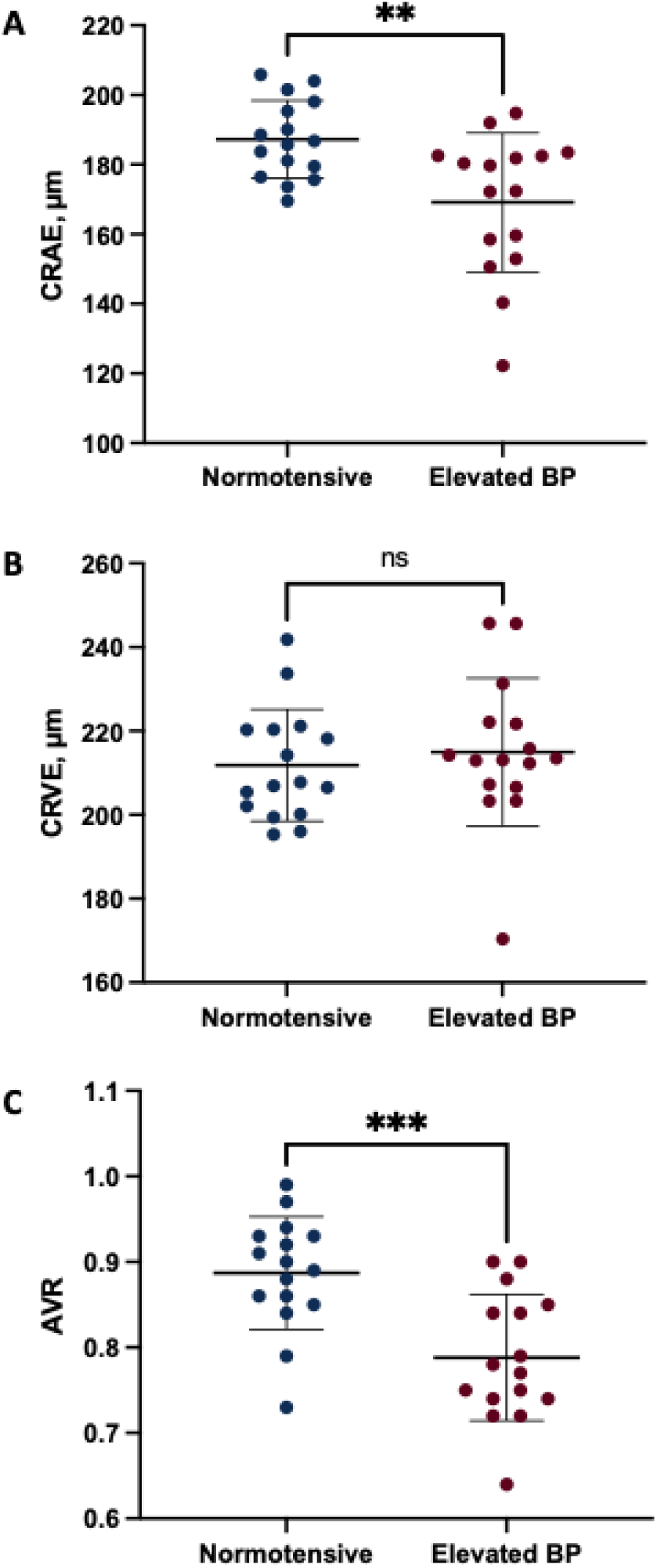
Retinal calibre assessment of subjects with normal versus elevated blood pressure (BP). Fundoscopy was used to assess **A,** central retinal arteriolar equivalent (CRAE), **B,** central retinal venular equivalent (CRVE), and **C,** arteriolar-to-venular-ratio (AVR). Comparison between normotensive and elevated BP groups by ANCOVA having sex as covariate (n=16 per group). Data presented as mean ± standard deviation (SD). ** P<0.01 and *** P<0.001.

### Association between endothelial cell function and retinal vessel diameters

We used linear regression analyses to assess correlations between all ECFC functional and microvascular parameters (complete regression data in **Supplemental Table S1**). Among all regressions adjusted for sex, we only observed significant negative correlations between the number of days taken for ECFC colony formation and CRAE (r^2^= 0.12, P=0.016, **Fig.3A**) and AVR (r^2^= 0.12, P=0.019, **Fig.3B**), with adjustment for glucose levels not changing the significance of these correlations.

**Figure 3.**
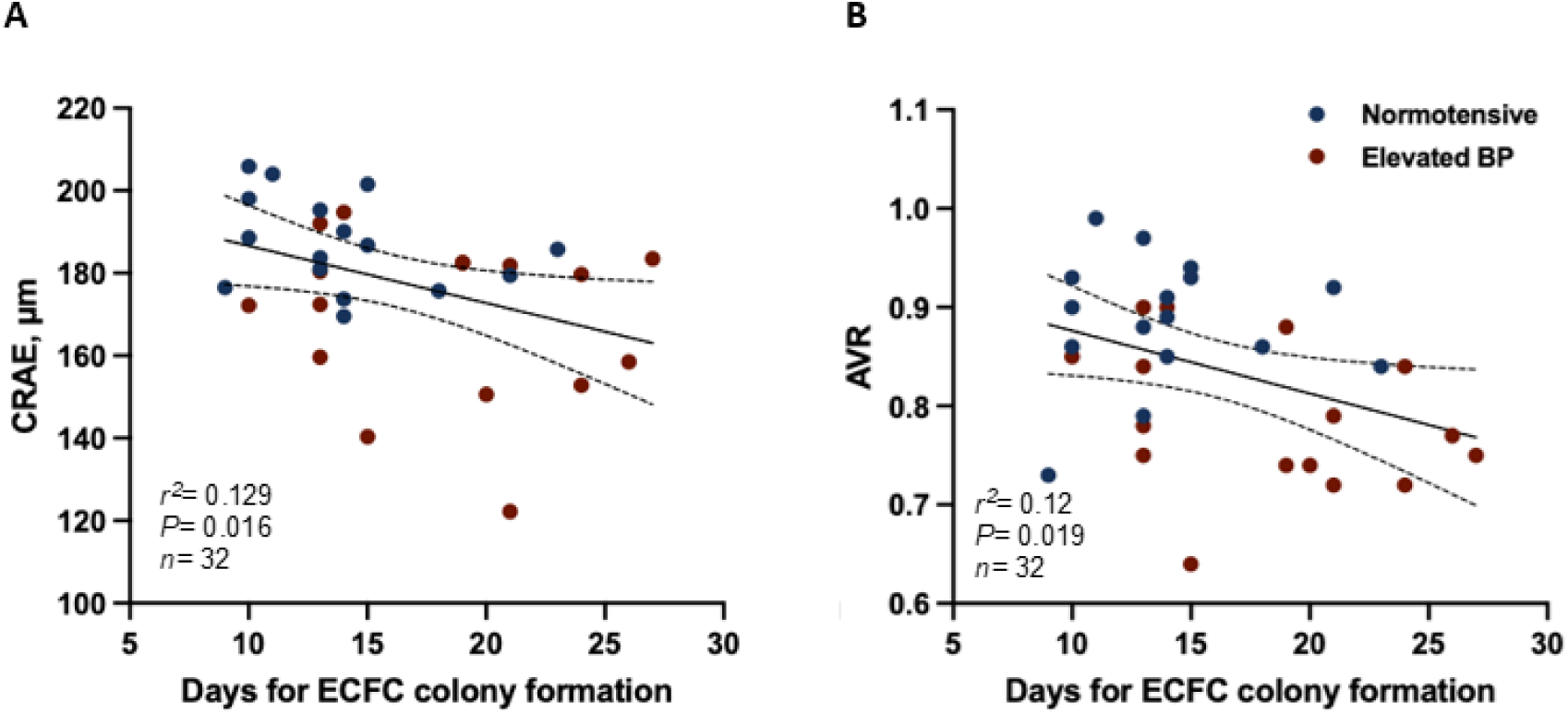
Linear regression between ECFC functional parameters and retinal vascular calibre. The days taken for ECFC colony formation is negatively correlated with **A,** central retinal arteriolar equivalent (CRAE) (r^2^= 0.185, P=0.016) and **B,** arteriolar-to-venular-ratio (AVR) (r^2^= 0.12, P=0.019). Linear regression adjusted for sex. Normotensive and elevated blood pressure (BP) individuals are represented by blue and red circles, respectively.

### Relationship between ECFC function and retinal vessel diameters with systolic BP

Complete linear regression data of systolic BP and ECFC function or microvascular phenotype are presented in **Supplemental Table S2** with significant results summarized in **Figure 4**. Negative correlations were observed only between systolic BP and ECFC proliferation rate (r^2^=0.155, p=0.036, **Fig.4A**), CRAE (r^2^= 0.505, P<0.001, **Fig.4B**) and with AVR (r^2^= 0.477, P<0.001, **Fig.4C**).

**Figure 4.**
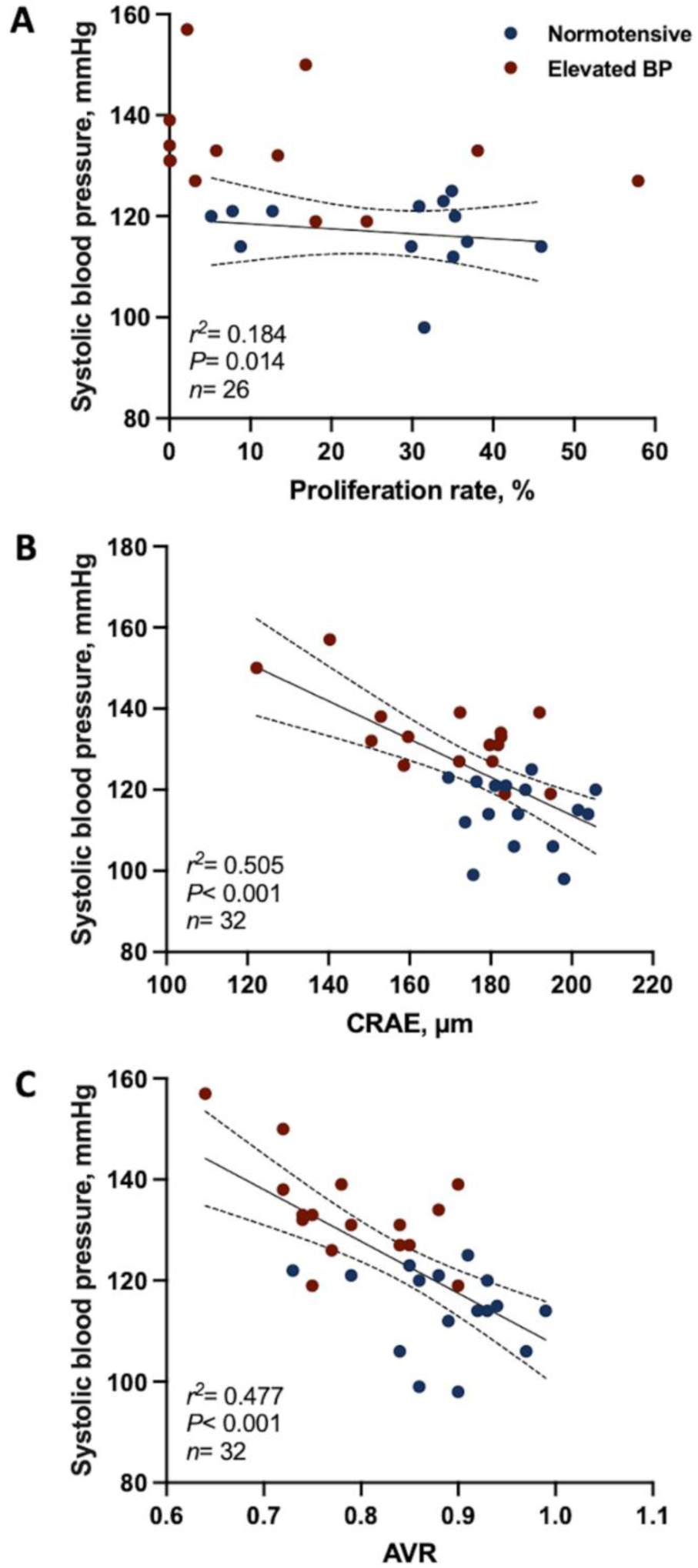
Relationship between systolic blood pressure (BP) and ECFC proliferation rate and retinal microvascular structure in young adults. Systolic BP negatively correlated with **A,** proliferation rate (r^2^= 0.155, n=26, P=0.036), **B,** central retinal arteriolar equivalent (CRAE) (r^2^= 0.505, n=32, P<0.001), and **C,** arteriolar-to-venular-ratio (AVR) (r^2^= 0.477, n=32, P<0.001). Linear regression adjusted for sex. Normotensive and elevated blood pressure (BP) individuals are represented by blue and red circles, respectively.

Multiple linear regression was used to estimate the relationship between ECFC function and retinal microvascular diameters with systolic BP in the study population (**Table 2**). ECFC proliferation rate and CRAE were included in the model along with participant’s sex and glucose levels as previous reports had demonstrated associations of sex- and glucose with ECFC function and retinal microvascular structure^34–36^. ECFC proliferation rate and CRAE had not correlated with each other, but had independently correlated with systolic BP (P<0.001). More specifically, a 0.37% decrease in systolic BP was observed for every 1% increase in ECFC proliferation rate (P=0.009) and a 0.75% decrease in systolic BP for every 1% increase in CRAE (P<0.001).

**Table 2.**
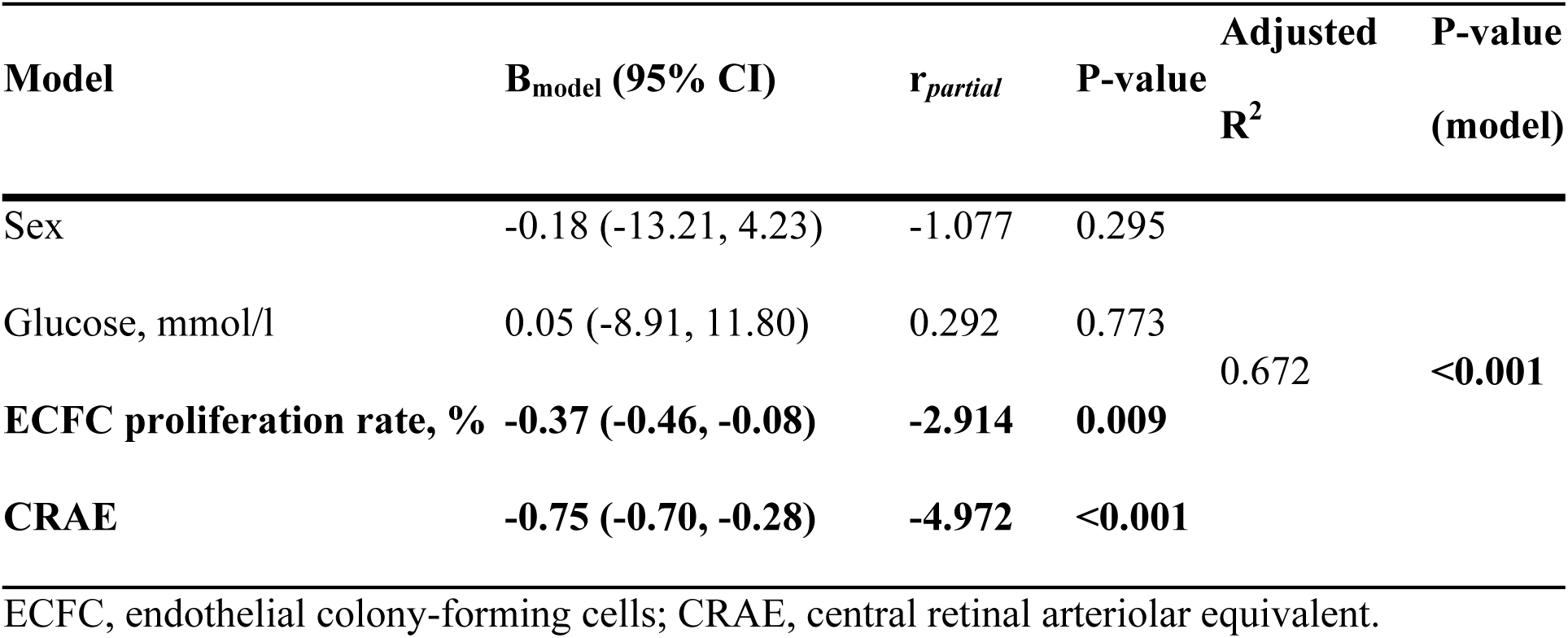
Multiple linear regression with systolic blood pressure (BP) as dependent variable.

### Proteomic signature of ECFCs from subjects with normal and elevated BP

The comparison of proteomic signatures of ECFCs identified 72 differentially expressed proteins between normal and elevated BP subjects, from a total of 2691 proteins, with unique peptides ≥2 and either fold change ≥1.5 or ≤-1.5 (67 upregulated and 5 downregulated proteins) **(Fig.5A)**. Heatmaps of n=72 identified proteins are shown in **Fig.5B**. String™ interactive pathway analysis of proteins further revealed two main interactive clusters (**Fig.5C**), one cluster containing mainly proteins related to extracellular matrix organisation (yellow circles) and blood coagulation (light blue circles), and a second cluster containing proteins related to regulation of coagulation (pink circles), exocytosis (purple circles) and haemostasis (green circles). The analysis of gene ontology revealed the top 10 enriched pathways being the following: 1) platelet degranulation, 2) regulated exocytosis, 3) regulation of haemostasis, 4) extracellular matrix organisation, 5) regulation of coagulation, 6) neutrophil mediated immunity, 7) blood coagulation, 8) secretion by cell, 9) neutrophil degranulation, and 10) negative regulation of wound healing **(Fig.5D)**. Heatmaps of proteins identified in each enriched pathway are shown in **Supplemental Figure S1**.

**Figure 5.**
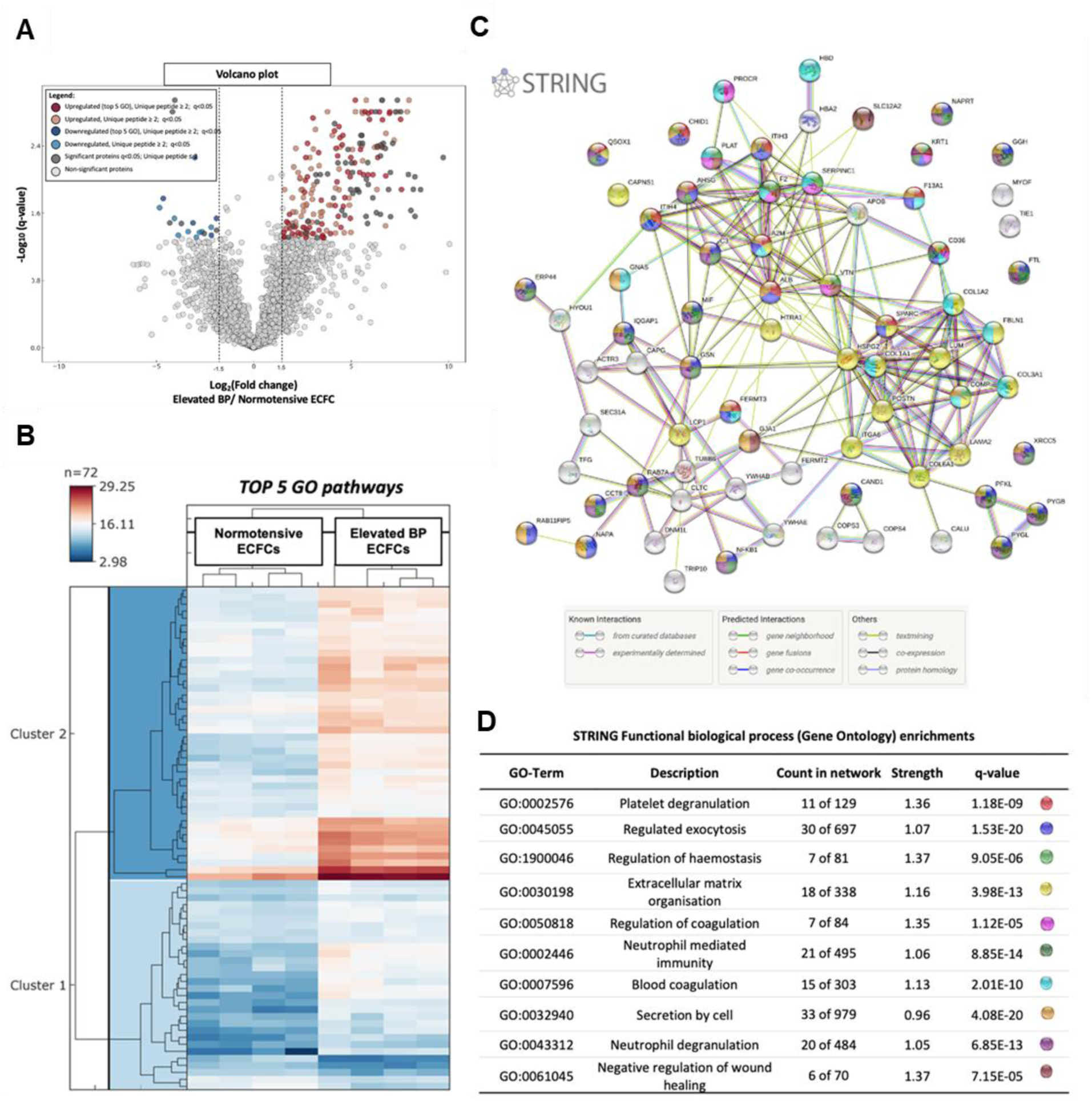
Proteins identified in top enriched gene ontology (GO) pathways differently expressed in ECFCs of subjects with normal versus elevated blood pressure (BP). A, Volcano plot shows significantly regulated proteins between normotensive and elevated blood pressure ECFCs, after FDR correction for multiple testing. A total of 2691 proteins were quantified with 158 proteins demonstrating significant differential abundance between the compared conditions based on unique peptides ≥ 2 and either log2 (fold change) ≥ 1.5 or ≤ -1.5 (145 upregulated and 13 downregulated proteins). Among the 150 proteins, 72 proteins were identified as part of top 5 GO pathways with 67 upregulated proteins and 5 down regulated proteins. **B,** Heatmaps of n=72 proteins involved in enriched GO pathways (n=72). Results are represented as a heatmap displaying protein expression levels on a logarithmic scale. Red indicates high expression while dark blue indicates low or no expression. **C,** String™ interactive pathway analysis and **D,** top 10 rank GO pathways identified with n=72 proteins.

We further identified 51 proteins of a total of 72 that significantly correlated with all four ECFC functional characteristics (**Supplemental Table S3**). Interaction diagram of identified 51 proteins indicate a potential central role of the growth factor receptor-bound protein-2 (GRB2) gene in regulating ECFC function in young adults with elevated BP (**Figure 6**). Further analysis of the identified 51 proteins revealed enrichments in GO biological pathways predominantly in vesicle mediated transport and exocytosis.

**Figure 6.**
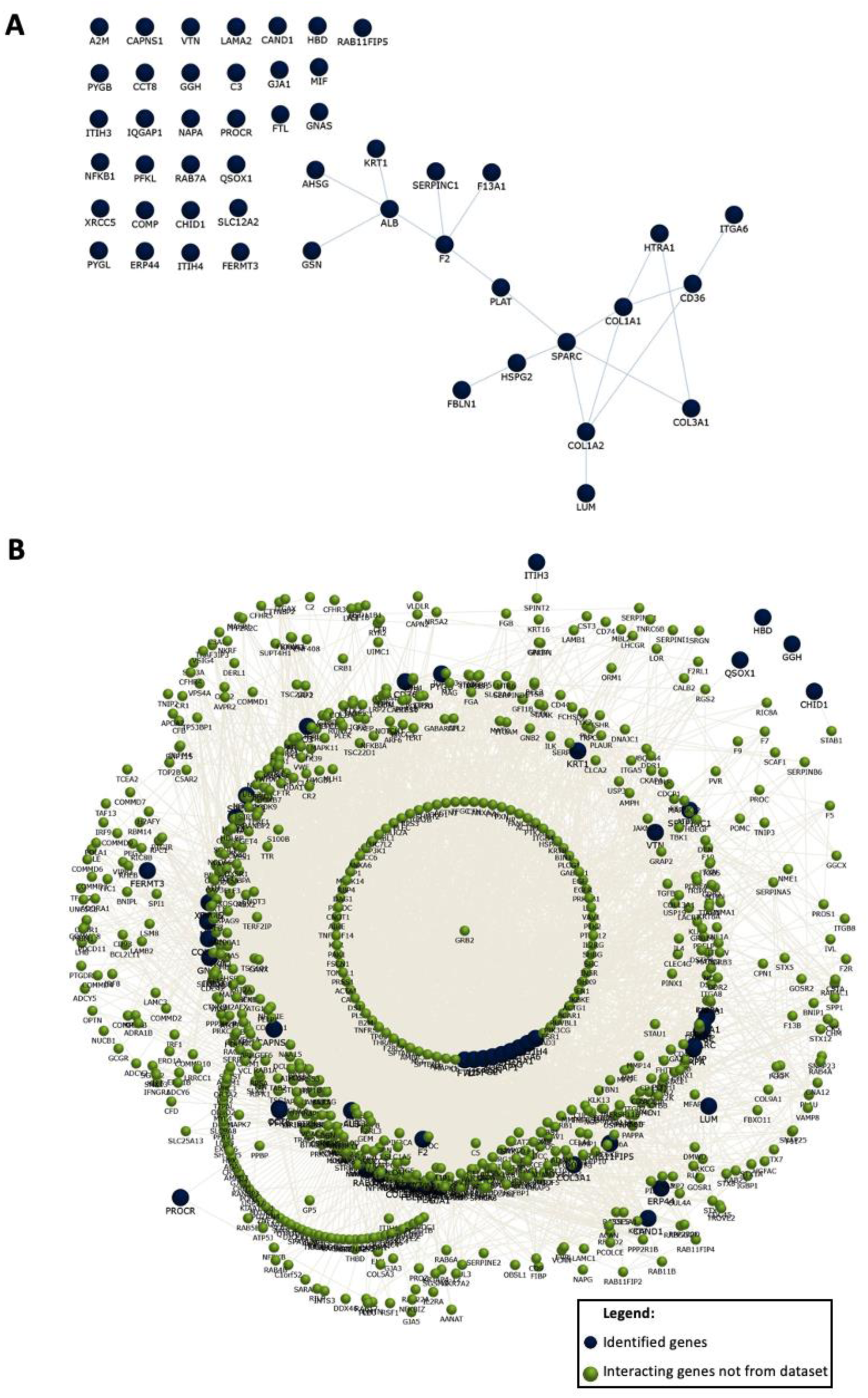
Interaction diagrams of n=51 identified proteins significantly correlated with ECFC function. **A**, Functional enrichment analysis revealed potential interactions between identified proteins and **B,** interactions between identified proteins (blue circles) with interacting genes/proteins not from dataset.

## DISCUSSION

Our results showed that young adults with elevated BP have a significant impairment in both expansion and vasculogenic potentials of ECFCs, which were associated with retinal microvascular impairments. Molecular evaluation of ECFCs underlined distinct proteomic profiles in cells from subjects with normal versus elevated BP with the activation of biological mechanisms related to extracellular matrix organization, coagulation, exocytosis and neutrophil mediated pro-inflammatory mechanisms in cells of subjects with elevated BP.

The association between retinal vascular diameters and hypertension has been well-established ^26, 34, 37^. Retinal arteriolar narrowing is being recognized as one of the earliest signs of hypertension and is inversely related to higher BP ^17, 38, 39^, which is in agreement with our findings. Furthermore, data from several longitudinal cohort studies provided the first prospective clinical evidence showing that narrower arteriolar diameters as reflected by a smaller CRAE and AVR not only relate to increased risk of incident hypertension but might precede the endothelial vascular dysfunction and clinical manifestation of CVD symptoms ^6, 7, 18, 39, 40^. These data, along with our data, suggests that early structural micro-arteriolar damage, visible in the retina as a sign of endothelial vascular dysfunction, may further precede the development and progression of hypertension as early as during adulthood.

In addition, for the first time, we showed the association between compromised endothelial cell function and narrowing of retinal arterioles that could potentially help to explain later manifestation and progression of hypertension. This has been observed in some studies revealing endothelial vascular dysfunction triggered by arteriolar narrowing and resulting shear-stress, leading to endothelium activation and inflammatory response ^41, 42^.

Under such conditions, the endothelium switches from a quiescent to an overactive state, where it starts to secrete vasoconstrictive factors, like endothelin-1 ^43^ and thromboxane ^44^, and proliferative factors, like VEGF (vascular endothelial growth factor), fibroblast growth factor 2 ^45^, CXCL12 ^46^, with reduced secretion of vasodilators, like nitric oxide (NO) and prostacyclin, indicating that endothelial and microvascular alterations might play a central role in the pathogenesis of hypertension ^41^.

Furthermore, our findings provided insights on potential mechanisms underlying vascular pathophysiology of hypertension in men and women, with endothelial and microvascular dysfunction potentially anticipating later BP changes in young-adult subjects. Endothelial vascular dysfunction is characterized by an abnormal remodelling of blood vessels stimulated by pro-thrombotic, pro-inflammatory, and pro-constrictive factors ^4^, which are mechanisms also activated during hypertension ^8^. It has been suggested that endothelial vascular dysfunction promotes arterial structural and functional abnormalities, including endothelial cell apoptosis or insufficient angiogenesis ^47^. Such responses have been described as secondary to a lower production of endothelial NO, which is a factor essential for capillary sprouting and maintenance of endovascular health ^48, 49^. Endovascular phenotype changes further contribute to promote vascular remodelling ^47, 49, 50^. Our molecular findings are in line with these biological pathways, showing the regulation of similar antiangiogenic mechanisms in dysfunctional circulating endothelial cells of subjects with elevated BP. Additionally, our findings that adverse indicators in the retinal microvasculature correlate to impaired ECFC function and antiangiogenic proteomic profiles offers further validation of the utility of fundoscopy for routing cardiovascular risk screening.

Our molecular approach also revealed GRB2 as a central protein potentially linking endothelial cell and microvascular dysfunction in subjects with elevated BP. GRB2 is known as an adaptor protein connecting growth factors with essential cellular responses driving cell survival and adaptive responses ^51^. Structurally, GRB2 is made of a SH2 domain, binding to phosphotyrosine motifs, and two SH3 domains, binding to prolin rich motifs ^51^. GRB2 also binds to the C-terminal part of Sos, an exchange factor of the small GTP-binding protein Ras, forming a complex that mediates Ras activation in response to various growth factors, including epidermal growth factor (EGF), VEGF and platelet-derived growth factor (PDGF). The complex is described to subsequently activate mitogen-activated protein kinases (MAPK), mainly ERK ^52^. This mechanism has been largely described in ERK1/2 activation by angiotensin II, being a critical molecular mechanism inducing vascular smooth muscle cells hypertrophic growth in models with high angiotensin II exposure ^53, 54^. GRB2 is therefore considered a complex molecule eliciting either pro-survival responses or promoting exacerbated maladaptive responses under pathological stress.

In endothelial cells, similar opposing effects have been described to GRB2, mainly depending on molecules binding to this protein. For example, Gab1 (GRB2-associated binder 1) binds to GRB2 after phosphorylation usually stimulated by VEGF, among other growth factors. This complex targets Akt, protein kinase A (PKA) and endothelial NO synthase (eNOS) in endothelial cells, or stimulate ERK1/2 and Kruppel-like factor 2 (KLF2) pathways ^55, 56^. Altogether, Gab1/GRB2 complex downstream pathways have been described to positively regulate angiogenesis by inhibiting apoptosis while stimulating proliferation, migration and tube formation ^56^. Contrastingly, Gab2, another GRB2 binding protein, has been described as a crucial molecule involved in the downstream signalling of pro-inflammatory interleukin receptors, antigen complexes, in addition to other growth factors ^51,57^. Pro-inflammatory effects of Gab2 has been associated with the activation of TAK1 (transforming growth factor beta-activated kinase 1) and p38 MAPK because Gab2 silencing can suppress the ubiquitination of TAK1 and activation of MAPKs and NFκB (nuclear factor kappa B), confirming an important role of Gab2 on cell inflammatory signalling and endothelial cell dysfunction ^57^.

To this date, GRB2 has not been directly associated with clinical hypertension, but it is associated with vascular remodelling in pulmonary arterial hypertension ^58^, diabetic retinopathy ^59^ and more broadly with postnatal angiogenesis ^58, 60, 61^. The role of this protein on postnatal angiogenesis and microvascular development has been systematically described using Gab1 and Gab2 knockout mice, which are viable mouse models without significant changes in vascular development. However, these models clearly develop changes in microvessel formation and endothelial cell function under stressful conditions. Endothelium-specific Gab1 knockout mice develop a remarkably impaired blood flow recovery and necrosis in the leg when submitted to hindlimb ischemia ^60^. Inversely, Gab2 knockout mice have suppressed expression of cell adhesion molecules in endothelial cells with reduced expression of inflammatory cytokines in response to various inflammatory stimuli, conferring vascular injury resistance in these mice ^57^.

In this study, we have shown that GRB2 was identified as a central protein integrating most of the 51 targeted proteins associated with impaired ECFC function and predominantly involved in mechanisms related to vesicle formation, mediated transport and exocytosis. A deeper molecular investigation of ECFC enriched proteins revealed a pro-inflammatory profile with platelet and neutrophil degranulation in dysfunctional cells. The immune effector response described in these cells further suggest an increased production and exocytosis of extracellular vesicles (EVs), which could be acting as inflammatory effectors in such responses. This is interesting, because EVs, either apoptotic bodies, micro-vesicles, or exosomes, have emerged as essential players in cell-cell communication in normal physiology and pathological conditions ^62^. EVs can carry ‘messages’ through their special cargoes, mainly consisting of cytokines, proteins, nucleic acids (mRNAs, and microRNAs), to recipient cells and lipids ^63–65^, also capable of modulating endothelial cell behaviour and angiogenesis ^65–69^. Furthermore, studies have shown that endothelial cell dysfunction can trigger the release of endothelial cell-derived EVs, and that these vesicles can play an increasing role in disease progression ^8, 14, 42, 70–72^. Increased concentration of endothelial cell- derived microparticles have been also shown to inhibit angiogenesis in hearts by inducing eNOS dysfunction ^73^. Recently, it was shown that endothelial cell-derived EVs can further inhibit tube-like structure formation *in vitro* and angiogenesis *in vivo* ^74^. Yet, the link between GRB2 and EVs formation or transport is still not completely understood. Our molecular findings suggest that changes in endothelial cell function may associate with factors regulating GRB2 that could ultimately lead to impaired angiogenesis and microvascular impairments in young-adults with elevated BP. Additional studies are needed to untangled these molecular pathways and their link with the vascular pathophysiology of hypertension.

A strength of our study was the ability to link up cellular properties with clinical measurements, including retinal microvasculature and BP, despite the limited sample size. We showed the interaction of endothelial cell function and retinal microvasculature and investigated their relationship with BP in men and women not receiving any antihypertensive medication. Furthermore, age and sex-matched individuals were used for our proteomic investigations to minimize any potential bias. One of the challenges faced in the study was the missing data mainly caused by limitations of ECFC expansion, particularly of dysfunctional cells ^2^, which have not allowed further detailed validation of targeted molecular mechanisms in dysfunctional endothelial cells.

In conclusion, we described for the first time the association between ECFC dysfunction and retinal arteriolar narrowing in young-adult men and women with elevated BP. Antiangiogenic mechanisms associated with cellular adaptations to growth factors involving the protein GRB2 may link endothelial cell dysfunction to impaired retinal microvascular structure in subjects with elevated BP. Further clinical and molecular studies are needed to untangle the role of GRB2, and key biological pathways regulated by this protein in endothelial and microvascular changes associated with hypertension.

## ACKNOWLEDGEMENTS

The authors thank the Oxford Cardiovascular Clinical Research Facility (CCRF) and the CCRF research nurses and the participants for their support, dedication, and contributions. The authors also thank Dr. Kessler’s laboratory at the Target Discovery Institute Mass Spectrometry, University of Oxford, for the mass spectrometry use and proteomic analyses.

## SOURCES OF FUNDING

This work was supported by Wellcome Trust (grant number 105741/Z/14/Z), the British Heart Foundation (grant number PG/17/13/32860), and the Oxford British Heart Foundation Centre of Research Excellence (grant number RE/13/1/30181). Dr Lewandowski is funded by a British Heart Foundation Intermediate Basic Science Research Fellowship (grant number FS/18/3/33292). Dr Bertagnolli is funded by grants from CIUSSS Nord-de-l’Île-de-Montréal, SickKids Foundation and Canadian Institutes of Health Research (grant number NI20-1037).

## DISCLOSURES

None

## SUPPLEMENTAL MATERIAL

Supplemental Tables S1–S3 Supplemental Figure S1

## Notes

### Competing Interest Statement

The authors have declared no competing interest.

